# Feature Importance in Predicting Clinical Outcome: Statistics vs. Explainable Artificial Intelligence

**DOI:** 10.1101/2024.07.21.604467

**Authors:** Parisa Amin

## Abstract

At the time of diagnosis for cancer patients, a wide array of data can be gathered, ranging from clinical information to multiple layers of omics data. Determining which of these data are most informative is crucial, not only for advancing biological understanding but also for clinical and economic considerations. This process facilitates the selection of the most significant markers, enhancing patient stratification and informing treatment recommendations. In this paper, we start with 89 features extracted from multiomics and clinical data and aim to identify the most important ones in predicting response to neoadjuvant chemotherapy (NAC) using different explainable Artificial Intelligence (XAI) models and statistics. Our results show that XAI methods consistently recover important features that are missed by statistics and vice versa, hinting towards the need for complementary implementation of these methods. Furthermore, we find that a myriad of features, from mutations to immune infiltration, affect the response to NAC in breast tumors.

## 1 Introduction

Feature importance is a critical concept in the field of data science and machine learning, referring to the process of identifying and ranking the most significant covariates in a dataset that influence the prediction outcome^1^. This concept is particularly relevant when dealing with cancer patients data where one has to handle complex and high-dimensional biological data collected at different scales^2^. Identifying the key features within cancer patients data can reveal crucial biomarkers and pathways associated with disease mechanisms, aiding in the development of targeted therapies and personalized medicine approaches^3,4^. Thus, feature importance not only streamlines the analytical process by focusing on the most impactful variables but also significantly contributes to advancing clinical research and improving patient outcomes^5^.

Feature importance evaluation encompasses a variety of methods ranging from statistical techniques to XAI approaches^6^. Traditional methods like LASSO (Least Absolute Shrinkage and Selection Operator)^7^ and t-tests^8^ have long been used to identify significant features by imposing regularization to prevent overfitting and by comparing means to determine statistical significance, respectively. These methods are robust and provide a solid foundation for feature selection. However, they may fall short when dealing with complex, non-linear relationships inherent in high-dimensional datasets. XAI methods, such as SHAP (SHapley Additive exPlanations)^9^ and LIME (Local Interpretable Model-agnostic Explanations)^10^, offer more detailed insights by interpreting model predictions and attributing importance scores to features. The necessity of performing both statistical and ML-based methods concurrently lies in the complementary insights they provide; statistical methods offer simplicity and statistical rigor, while modern methods enhance interpretability and account for intricate patterns in the data^11^. Comparing results from both approaches provides a comprehensive map of feature importance a better understanding of feature significance, leading to more robust and reliable conclusions in data analysis^12^.

In this paper, we first implement three statistical models: t-test, L1, and elastic net (EN) LASSO^13^ to explore the association of covariates with pathological complete response (pCR)—that is, no invasive and no in situ residuals in breast and nodes after treatment^14^. After performing the statistical analysis, we implemented several ML models to predict pCR (vs. residual disease, RD). To assess feature importance across these ML models, we employed the SHAP method. For each ML model, we analyzed the SHAP values to identify the most important features. Finally, we compared the results of the ML models with each other and with the statistical methods. This comparative analysis provided a holistic view of feature importance, highlighting both consistent findings and unique insights offered by each approach. The integration of statistical methods and XAI techniques provided a robust framework for feature importance, enhancing our understanding of the key predictors of pCR.

## 2 Results

We have a dataset where a wide range of features were extracted for 149 patients^15^, which includes a variety of omics data as well as pathological and clinical data. **Tumor features**: sWES (shallow whole exome sequencing) and WGS (whole genome sequencing) data were used by^15^ to extract such as chromosomal instability, purity and ploidy using ASCAT (Allele-Specific Copy number Analysis of Tumors)^16^ as well as homologous recombination deficiency (HRD) and 30 mutational signatures^17^. These signatures help in understanding the underlying mutational processes and genetic alterations driving tumorigenesis. **Tumor Immune Infiltration**: Extracted from bulk RNA seq data, this category includes measures such as the Immunophenoscore^18^ and MCPcounter^19^. The Immunophenoscore assesses the immune-related components within the tumor microenvironment, while MCPcounter quantifies the abundance of various immune and stromal cell populations. **Cell Fractions**: Based on digital pathology data, fractions of three main cell types within the tumor microenvironment, including cancer cells, lymphocytes, and stromal cells were extracted. This information provides a detailed view of the cellular composition of the tumors. **Clinical Features**: This category encompasses crucial clinical characteristics such as Estrogen Receptor (ER) status, Human Epidermal Growth Factor Receptor 2 (HER2) status, PAM50 molecular subtype, tumor size, stage, lymph node status, and treatment regimen. These features are essential for understanding the clinical context and therapeutic responses of the patients. Overall, we included 89 of these previously extracted features in our analysis.

### 2.1 statistical methods

#### t-test

For each feature in our dataset, we performed an independent t-test to assess whether there was a statistically significant difference between the pCR and RD groups. Given the large number of features tested, multiple comparisons can increase the likelihood of false-positive results. To mitigate this risk, we applied the Bonferroni correction to adjust the p-values using rstatix^20^ R package. This correction method reduces the chance of type I errors by adjusting the significance threshold according to the number of tests conducted, providing a more reliable indication of significant differences between the two response groups.

For simplicity and clarity, we visualized the results by plotting the transformed adjusted p-values. Specifically, we plotted “-Log(p.adj)*Direction” for each feature, where “Direction” indicates the sign of the mean difference between the groups. This transformation not only emphasizes significant differences but also indicates the direction of the effect (positive or negative). Figure 1 illustrates these results, highlighting the features that show significant differences across the pCR and RD groups.

**Figure 1.**
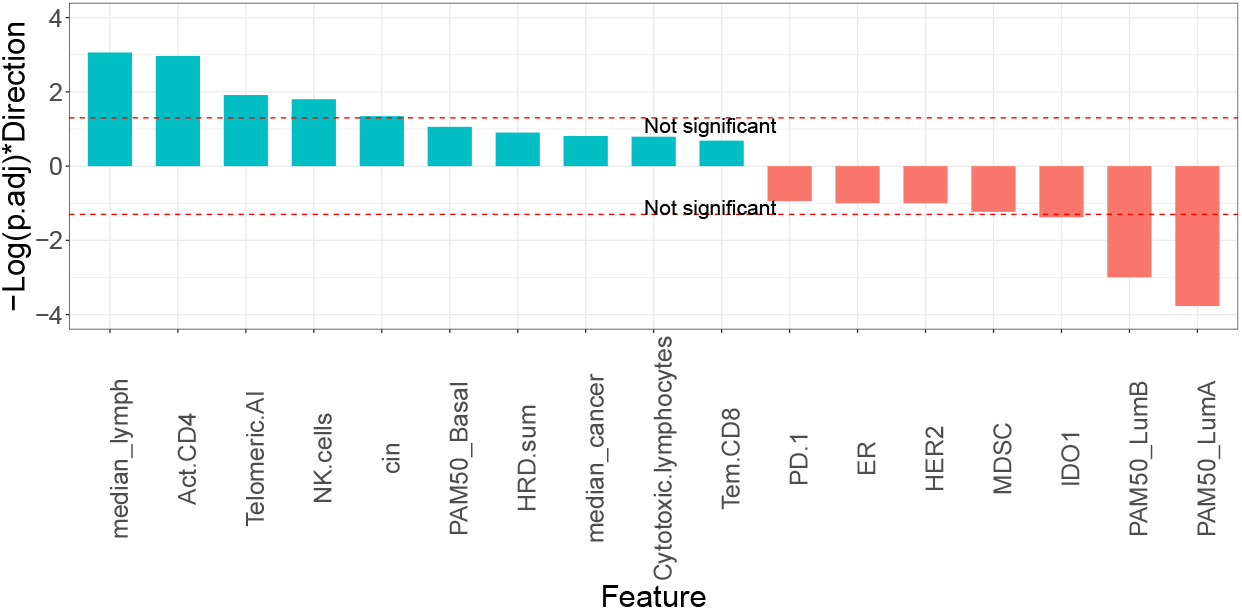
Comparison of feature differences between two response groups (pCR vs RD). The plot displays the transformed adjusted p-values using the formula “-Log(p.adj)*Direction” for each feature. Features with significant differences are highlighted, indicating their importance in distinguishing between the pCR and RD groups. Positive values suggest higher feature values in the pCR group, while negative values indicate higher values in the RD group.

#### LASSO

As a complementary approach to identify the most important features, we also implemented LASSO with L1 regularization. LASSO is particularly effective when aiming to enhance the prediction accuracy and interpretability of the statistical model it generates. By imposing a constraint on the sum of the absolute values of the model parameters, LASSO effectively performs feature selection by shrinking some coefficients (*β*s) to zero. This allows us to identify a subset of features that are most predictive of the outcome variable. We used the glmnet R package^21^ to run LASSO. Figure 2 (a) shows the Mean-Squared Error vs. *log*(*λ*). Sorted non-zero *β*s for the selected model with the best performance, judged based on Mean-Squared Error, are shown in Figure 2 (b). Similarly, we ran Elastic Net (EN) LASSO (L2 LASSO identifies all features as important). Figure 2 (c) shows the Mean-Squared Error vs. *log*(*λ*) for EN with *α* = 0.2. Sorted non-zero *β*s for EN LASSO are shown in Figure 2 (d).

**Figure 2.**
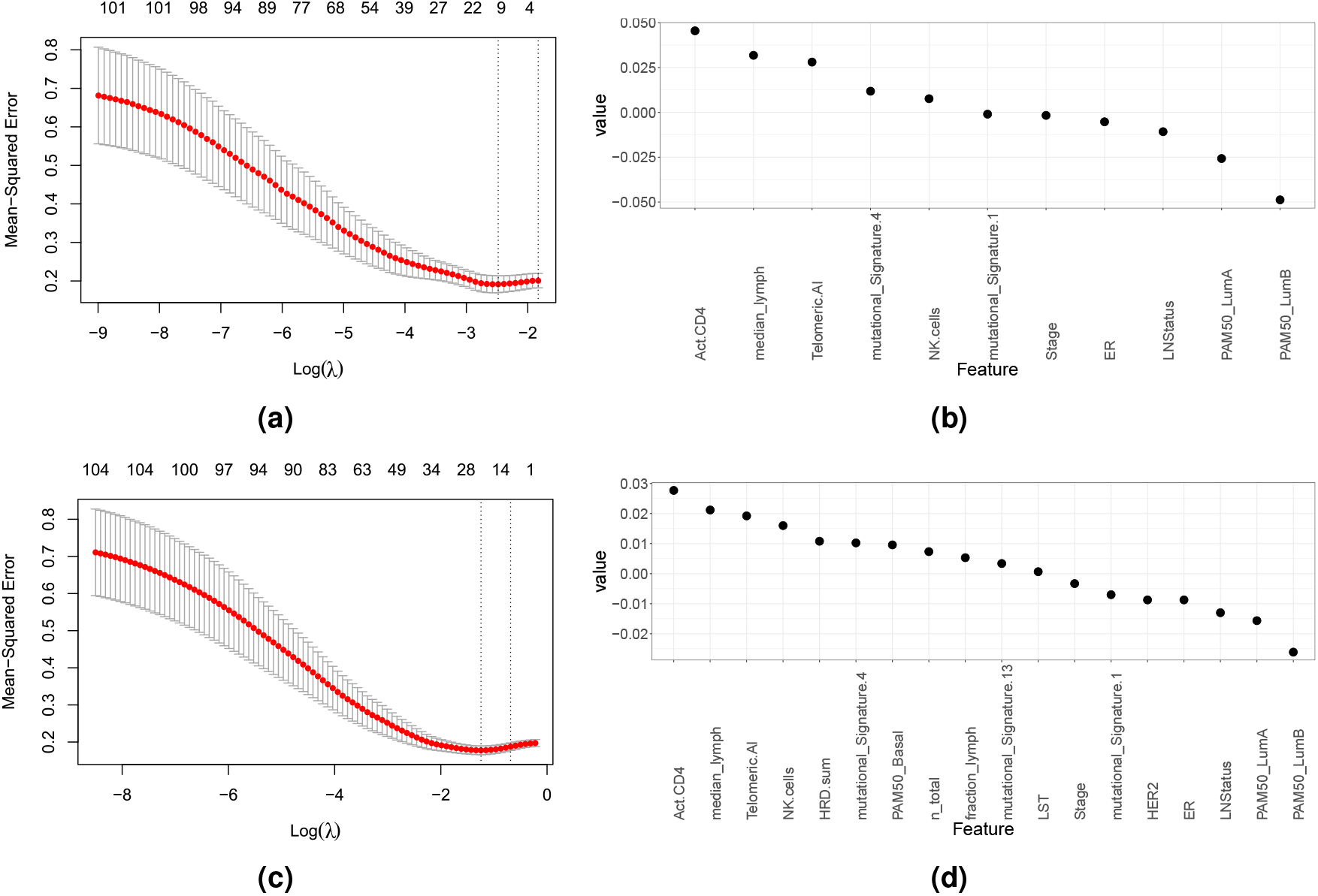
Feature selection using LASSO regression. (a) Mean-Squared Error in L1 LASSO vs *log*(*λ*). (b) top 11 features sorted by their coefficients, illustrating the importance of each feature in the model. Positive coefficients indicate features with a positive association with pCR, while negative coefficients indicate a negative association with it. This visualization highlights the most influential features selected by LASSO, providing insights into their relative importance in predicting pCR. (c) EN Model performance as a function of the number of features included. (d) Features with non-zero *β*s in EN.

### 2.2 ML models

After performing the statistical analysis, we implement machine learning models as follows:

#### Support Vector Machines (SVMs)

We employed SVMs to predict pCR using all 89 features in our dataset. The data was split into training and test sets with a ratio of 67% for training and 33% for testing to ensure robust model evaluation. To optimize the SVMs model, we performed used GridSearchCV where we searched through a predefined hyperparameter space to identify the best combination of parameters that maximize the model’s performance based on Area under curve (AUC) in receiver operating characteristic (ROC) graph. The best model has the following performance on 5-fold cross-validation: Accuracy: 70.529% (±6.612), f1 score: 0.564 (±0.078) and AUC: 0.808 (±0.077). The optimal SVMs model was fitted to the entire dataset and then Kernel explainer was used to calculate SHAP values. The top 20 features identified by SHAP, which have highest average absolute SHAP values (average |SHAP values|), are plotted in Figure 3.

**Figure 3.**
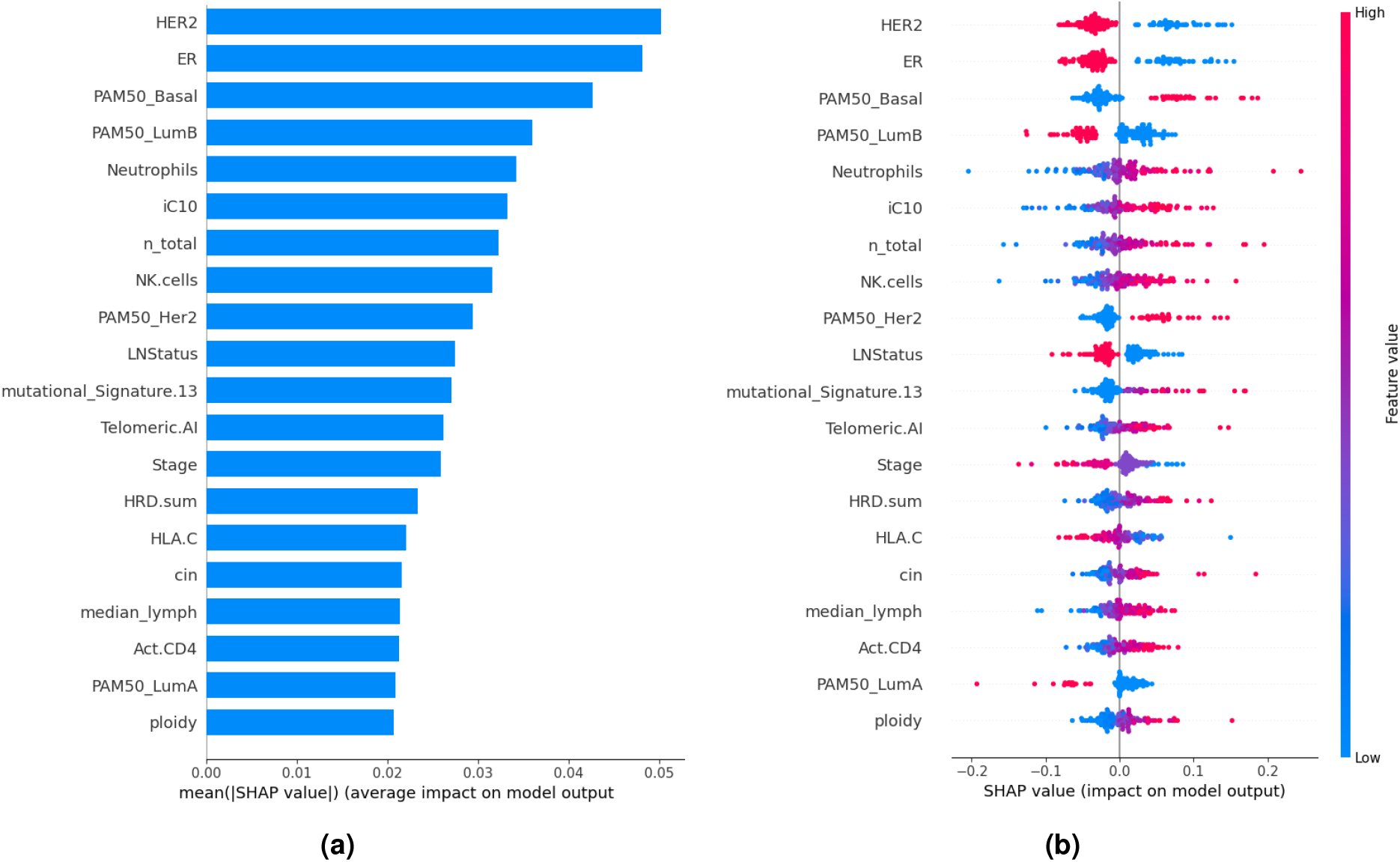
Feature importance analysis using SHAP for the Support Vector Machine (SVM) model. Panel (a) displays the top 20 features ranked by their SHAP values, indicating the magnitude of each feature’s contribution to the model’s prediction of pCR. Panel (b) illustrates the direction of the impact for each feature, where positive SHAP values suggest a positive influence on predicting pCR, and negative SHAP values indicate a negative influence. This visualization provides an understanding of the most influential features and their respective effects on the model’s output.

#### Logistic Regression (LR)

LR was implemented to predict pCR using all 89 features in our dataset. GridSearchCV was used to identify the best parameters that enhance model performance based on AUC. The best model has the following performance on 5-fold cross-validation: Accuracy: 73.2 % (±9.1), f1 score: 0.635 (±0.085) and AUC: 0.813 (±0.068). This model was fitted to the entire dataset and Kernel explainer was used to calculate SHAP values. Top 20 features in LR model are shown in Figure 4 (a).

**Figure 4.**
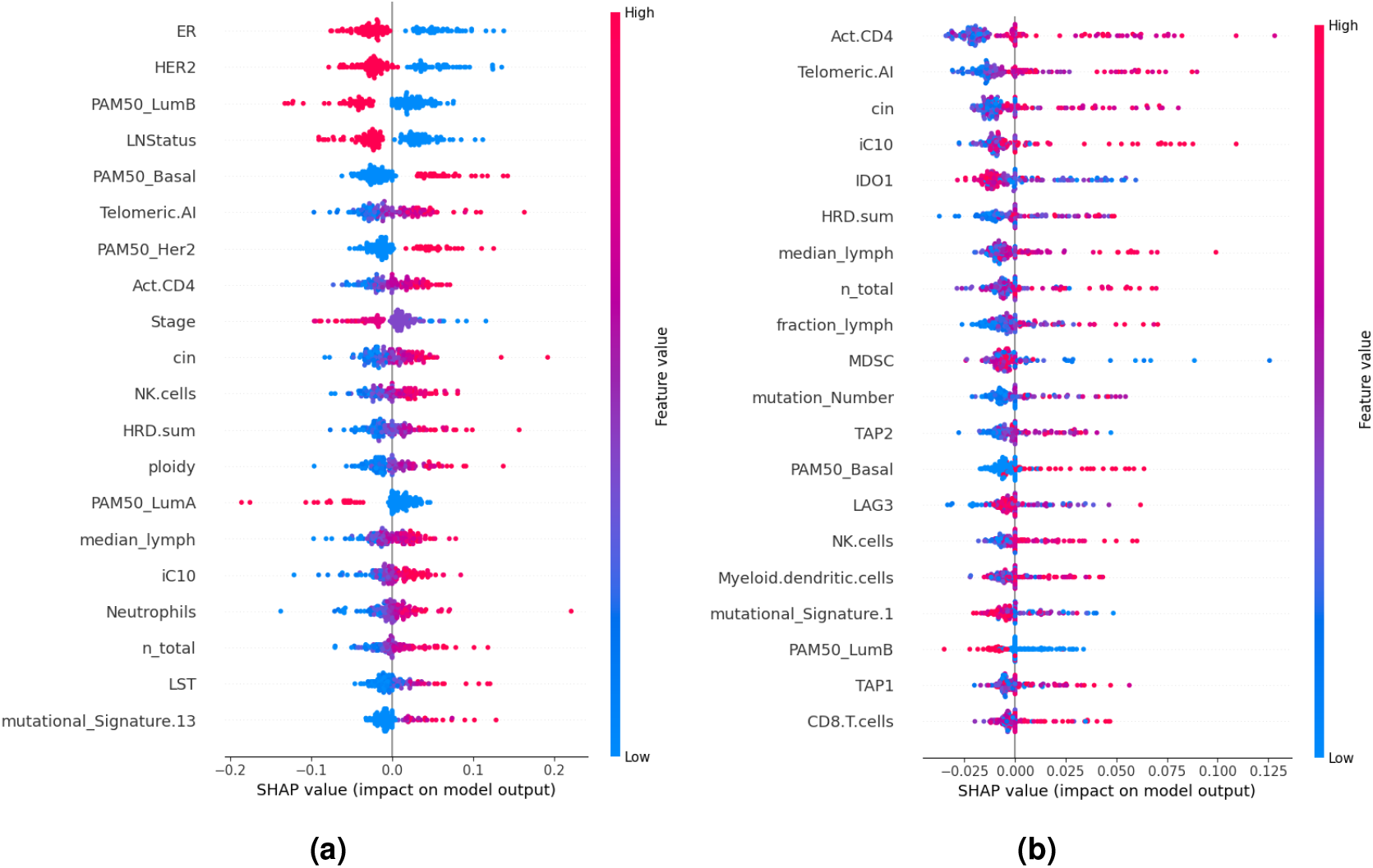
Top 20 features with the direction of the impact for each feature using LR (a) and RF (b). Positive SHAP values suggest a positive influence on predicting pCR, and negative SHAP values indicate a negative influence.

#### Random Forests (RF)

We employed the RF model to predict pCR. To optimize the RF model, we performed hyperparameter tuning using GridSearchCV. Similar to previous models, the objective was to identify the best combination of parameters that maximizes the model’s performance based on AUC. The best model has the following performance: Accuracy: 70.483 % (±8.222), f1 score: 0.302 (±0.102) and AUC: 0.750 (±0.082). This model was fitted to the entire dataset and SHAP values were calculated using Kernel Explainer. Top 20 features in RF are shown in Figure 4 (b).

#### Statistics vs XAI

To compare the XAI methods across different models, we first plotted the top 20 features for each model based on their SHAP values. This visual representation allowed us to directly observe and compare the most influential features identified by SVMs, LR, and RF. By displaying these features side-by-side, as shown in Figure 5 (a), we assess the consistency and divergence in feature importance rankings across models. This approach provided a clear overview of which features were deemed critical by each model, highlighting both common and unique patterns in how the models interpret the data.

**Figure 5.**
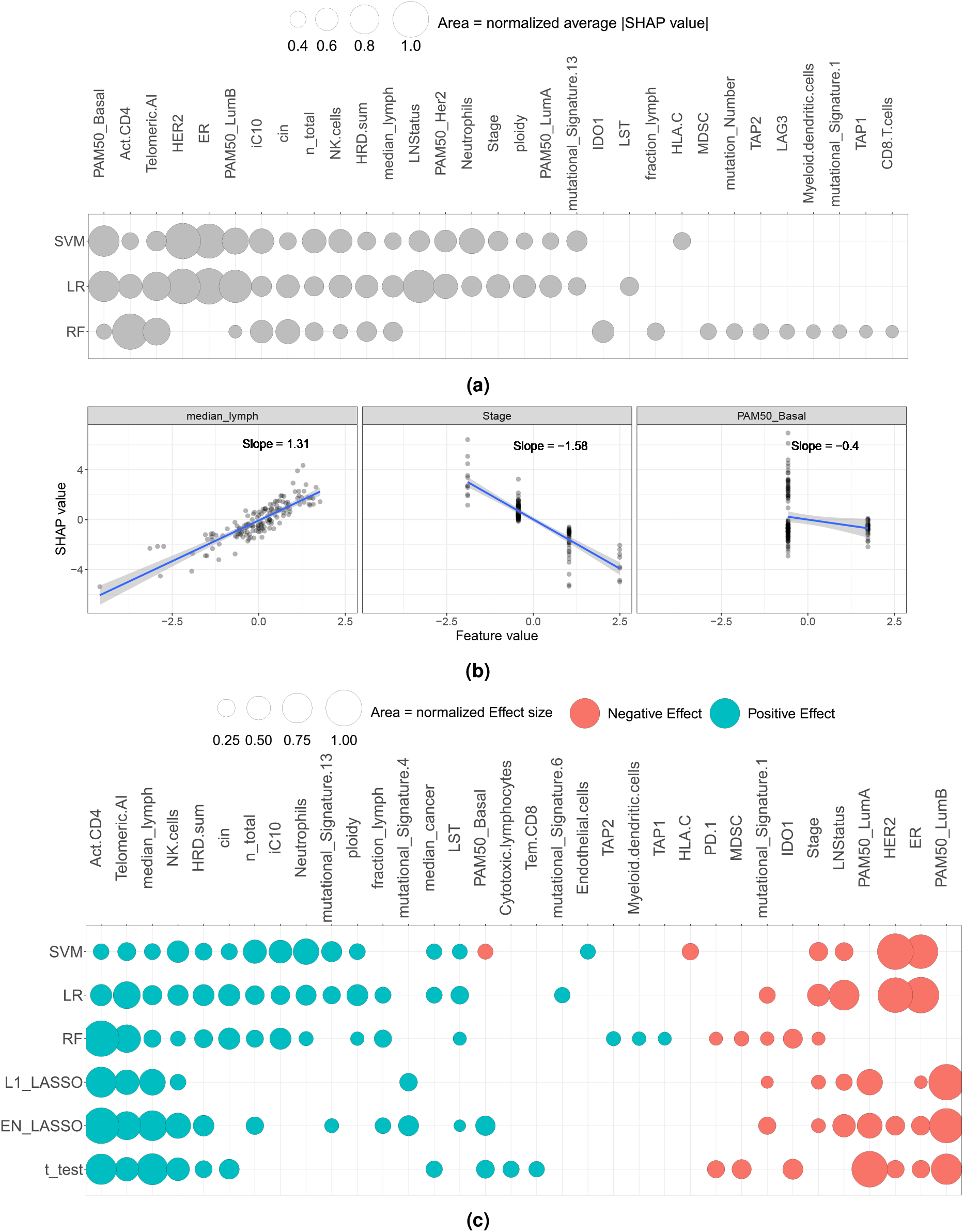
(a) Top 20 features identified by SHAP values across three different models: SVMs, LR, and RF. This visualization highlights the commonalities and differences in feature importance rankings among the models. (b) SHAP value vs. normalized feature values for three features in the LR model. These slopes were then normalized by dividing by the largest absolute value slope, yielding the “Effect”. Having a larger average |SHAP values| does not directly lead to a larger Effect, as PAM50–Basal shows a smaller value for slope and then Effect despite having larger |SHAP values| than median–lymph and Stage. (c) Comparison of the top 20 features with the most significant Effect, alongside the normalized outputs from statistical methods: L1 LASSO, EN LASSO, and t-tests.

Furthermore, to compare the effect of each feature in a more detailed manner, we analyzed the SHAP values by fitting a line to the SHAP value vs. normalized feature values for each feature. Following that, we normalized the slopes of these lines to quantify the strength of each feature’s effect on the model’s predictions, and named this measure “Effect”. It is worth mentioning that these Effects capture the role of features in model predictions and, from a biological perspective, reflect the association between the features and clinical outcome (pCR). This measure is different from the average |SHAP values| as it directly accounts for the directionality (positive or negative) of feature contributions in model predictions. Figure 5 (b) shows the slope for SHAP values vs. normalized feature values for three different features in the LR model. In this case, PAM50*_*Basal, while having a larger average |SHAP values| compared to both median*_*lymph and Stage, shows a smaller slope, revealing a less direct effect in model predictions.

By selecting the top 20 features with the most significant Effects based on these normalized slopes, we were able to identify the most impactful variables. For comparability, we also normalized the outputs from statistical methods (*β*s in L1 and EN LASSO and -Log(p.adj)*Direction in t-test) and used the same naming convention. Subsequently, ML top features were compared against the results obtained from statistical methods, as shown in Figure 5 (c). This comparison allowed us to evaluate the consistency and discrepancies between modern XAI techniques and statistical approaches, providing an understanding of feature importance across different analytical methodologies.

## 3 Discussion

When dealing with multiple covariates, it is of great interest to identify the most important and informative ones. In the case of biological data, this necessity is even more pronounced. This is partly due to the need and emphasis on predictions and classification in biological science and the concept of biomarkers^22^.

In our analysis, we employed a multi-faceted approach to feature importance evaluation by utilizing statistical methods and XAI techniques. Specifically, we used LASSO L1, EN LASSO, and t-tests for traditional feature selection. In parallel, we applied SHAP to evaluate feature importance in three different ML models.

Interestingly, our results revealed notable similarities in the key features identified across the different models and methods. Certain features consistently emerged as significant, underscoring their robust influence on the clinical outcome. However, we also observed differences in the feature importance rankings among different approaches. These discrepancies highlight the unique ways in which each model and method captures and interprets the data. For instance, RF, with its ability to model complex interactions, sometimes identified different sets of important features compared to linear models like LR and SVMs. Furthermore, while LASSO and EN LASSO focused on sparsity and variable selection through regularization, SHAP provided a more detailed interpretation by attributing importance scores based on the contributions of each feature to individual predictions.

XAI and statistical methods identify their own unique set of features. In other words, SHAP results for different models are more similar than that of the statistical methods. This similarity between different ML models is even more apparent in comparison of SVMs and LR, as Figure 5 (a) shows. Such similarity likely stems from similarities in model structures in LR and SVMs as they both take a probabilistic model and minimizing some cost associated with misclassification based on the likelihood ratio^23^.

These findings underscore the value of employing a diverse array of feature selection methods and models. By comparing the results, we obtain a more holistic view of feature importance, ensuring that critical variables are not overlooked due to the limitations inherent in any single method or model. This integrated approach is particularly crucial in the context of multiomics data, where the complexity and high dimensionality necessitate robust and multifaceted analytical strategies. Moreover, these findings highlight the inherent limitations of previous attempts to combine statistics and ML methods for feature importance^15^. Implementing a two-step feature importance evaluation method may result in the loss of features not identified as important by statistical methods, which might have been recognized as significant if evaluated by ML methods.

From a biological perspective, different methods consistently identify three features as having a positive effect on pCR: infiltration of activated CD4 T cells (Act. CD4), median lymphocyte density in H&E images (median*_*lymph), and infiltration of natural killer (NK) cells. All three features represent immune infiltration in the TME. Our finding on the positive role of median–lymph in response to NAC agrees with previous results^24^. Further research is required to clarify the relationship between median*_*lymph and NK and Act. CD4 T cells. In particular, to explore whether these features reflect the same biological process at different levels (bulk RNA vs. H&E) or are independent processes. Such an analysis, at least in theory, could clarify if gathering only one type of data (bulk RNA-seq or H&E) is informative enough.

Additionally, telomeric allelic imbalances (AI) are the number of AIs (the unequal contribution of parental allele sequences with or without changes in the overall copy number of the region) that extend to the telomeric end of a chromosome^25^. Our findings here are in line with previous results on the positive association between telomeric AI and response to NAC in breast cancer^26^.

Furthermore, our findings highlight that a myriad of features affect clinical outcomes. A combination of immune infiltration, mutation signatures and clinical features, such as ER status, influences patient prognoses. This observation exhibits how genetic alterations, biological markers, and clinical characteristics converge to affect clinical outcomes. This holistic understanding emphasizes the importance of multi-dimensional approaches in biomedical research and the necessity of implementing complementary analytical methods to enable more accurate predictions and personalized treatment strategies.

## 4 Methods

### 4.1 ML Models

In this study, we implemented three machine learning models: SVMs, LR, and RF. Models were developed by using the Scikit-Learn library^27^ within the Python programming language. Hyperparameter tuning was performed using GridSearchCV, with the objective of maximizing AUC in all models.

#### 4.1.1 SVMs

The hyperparameters, including the regularization parameter *C*, kernels, and the kernel coefficient *γ*, were optimized using grid search with cross-validation to prevent overfitting and ensure the generalizability of the model.

#### 4.1.2 LR

LR, a linear model, was used due to its simplicity and interpretability in predicting binary outcomes. The regularization strength *C* was determined through grid search with cross-validation.

#### 4.1.3 RF

RF, an ensemble learning method, was employed to leverage its ability to handle high-dimensional data and complex interactions between features. The hyperparameters, including the number of estimators, maximum depth, and number of leaves were optimized using GridSearchCV.

### 4.2 SHAP Values

To elucidate the contribution of individual features, we employed SHAP^9^, a post-hoc explanation method that quantifies the marginal contribution of each feature to the prediction for each sample. This method is grounded in cooperative game theory and ensures consistency and local accuracy, making it an ideal tool for interpreting complex machine learning models. By using SHAP values, we were able to gain detailed insights into how each feature influenced the model’s predictions. This analysis was crucial for identifying the most impactful features that drive prediction of clinical outcomes. For all models, SHAP values were calculated using Kernel Explainer.

### 4.3 Statistics

In addition to ML models, we applied several statistical methods to evaluate feature importance, utilizing R for implementation. The methods employed were t-tests, L1 LASSO, and EN LASSO, with the following specifics:

#### 4.3.1 t-test

We performed t-tests using rstatix^20^ R package to identify features that have a statistically significant difference in means between the two outcome groups.

#### 4.3.2 L1 LASSO

LASSO regression with L1 regularization was used to perform feature selection and regularization simultaneously. This method is effective in high-dimensional settings as it tends to shrink coefficients of less important features to zero, thus performing variable selection. The optimal regularization parameter *λ* was determined using cross-validation in glmnet^21^ R package.

#### 4.3.3 EN LASSO

We also used EN LASSO, which combines both L1 and L2 regularization penalties, controlled by the mixing parameter *α*. For our analysis, we set *α* = 0.2 to balance between the two types of regularization. This approach is particularly useful when dealing with correlated features, as it can select groups of related features together. The regularization parameters were optimized through cross-validation in glmnet^21^ R package.

### 4.4 Data

Clinical variables and features extracted from the multiomics dataset were obtained from the publicly available repository at https://github.com/cclab-brca/neoadjuvant-therapy-response-predictor/tree/master/data. This dataset integrates various omics data types, including genomics, RNA seq, and clinical features, providing a view of the biological and clinical factors influencing NAC response in breast cancer. The rich multi-layered data allowed us to perform a detailed analysis of feature importance, leveraging both statistical methods and modern XAI techniques to identify key determinants of clinical outcomes.

### 4.5 Code Availability

Codes for this paper are available at https://github.com/ParissaAmin.

## Notes

https://github.com/cclab-brca/neoadjuvant-therapy-response-predictor/tree/master/data

### Competing Interest Statement

The authors have declared no competing interest.

